# Positive association between *Brucella* spp seroprevalences in livestock and humans from a cross-sectional study in Garissa and Tana River Counties, Kenya

**DOI:** 10.1101/658864

**Authors:** S.W. Kairu-Wanyoike, D. Nyamwaya, M. Wainaina, J. Lindahl, E. Ontiri, S. Bukachi, I. Njeru, J. Karanja, R. sang, D. Grace, B. Bett

## Abstract

**Background:** *Brucella* spp. is a zoonotic bacterial agent of high public health and socio-economic importance. It infects many species of animals including wildlife, and people may get exposed through direct contact with an infected animal or consumption of raw or undercooked animal products. We implemented a linked livestock-human cross-sectional study to determine seroprevalences and risk factors of *Brucella* spp. in livestock and humans. We also estimated intra-cluster correlation coefficients (ICCs) for these observations at the household and village levels.

**Methodology:** The study was implemented in Garissa County (specifically Ijara and Sangailu areas) and Tana River (Bura and Hola) counties. A household was the unit of analysis and the sample size was derived using the standard procedures. Serum samples were obtained from selected livestock and people from randomly selected households. Humans were sampled in both counties while livestock could be sampled only in Tana River County. Samples obtained were screened for anti-*Brucella* IgG antibodies using ELISA kits. Data were analyzed using generalized linear mixed effects logistic regression models with the household (herd) and village being used as random effects.

**Results:** The overall seroprevalences were 3.47% (95% confidence interval [CI]: 2.72 – 4.36%) and 35.81% (95% CI: 32.87 – 38.84) in livestock and humans, respectively. In livestock, older animals and those sampled in Hola had significantly higher seroprevalences that younger ones or those sampled in Bura. Herd and village random effects were significant and ICC estimates associated with these variables were 0.40 (95% CI: 0.22 – 0.60) and 0.24 (95% CI: 0.08 – 0.52), respectively. For human data, older people, males, and people who lived in pastoral areas had significantly higher *Brucella* spp. seroprevalences than younger ones, females or those who lived in irrigated or riverine areas. People from households that had at least one seropositive animal were 3.35 (95% CI: 1.51 – 7.41) times more likely to be seropositive compared to those that did not. Human exposures significantly clustered at the household level; the ICC estimate obtained was 0.21 (95% CI: 0.06 – 0.52).

**Conclusion:** The presence of a seropositive animal in a household significantly increased the risk of exposure in people in that household. *Brucella* spp. exposures in both livestock and humans clustered significantly at the household level. This suggests that risk-based surveillance measures, guided by locations of primary cases reported, either in humans or livestock, can be used to detect *Brucella* spp. infections livestock or humans, respectively.

**Author summary:** Brucellosis is an important zoonotic disease that primarily affects livestock and wildlife. In humans, the disease is characterized by prolonged fever, body aches, joint pains and weakness while in livestock, the disease causes abortions and infertility. We carried out a study in northeastern Kenya (Garissa and Tana River Counties) to identify factors that affect the distribution of the disease in people and livestock. Livestock and people from randomly selected households were recruited and serum samples obtained for screening using anti-*Brucella* IgG ELISA kits to determine their *Brucella* spp. exposure. Data obtained were analyzed using mixed effects logistic regression models. Results obtained show that human and animal *Brucella* spp. seroprevalences cluster at the household level. The odds of exposure in humans were at least three times higher in households that had at least one seropositive animal compared to those that did not. These results can be used to design risk-based surveillance systems where each *Brucella* infection identified in livestock or humans could signal potential locations of other secondary infections.

## Introduction

Brucellosis is a zoonotic disease caused by gram-negative intracellular coccobacilli of the family *Brucellaceae.* It is an economically important disease characterized by prolonged fever, night sweats, body aches, arthralgia and weakness in humans and by abortions and infertility in livestock [1]. There are at least ten *Brucella* spp.; six of these, i.e., *B. melitensis, B. abortus, B. suis, B. canis, B. ovis and B. neotomae* are considered classical species [2], with the first four being pathogenic to man [3]. *Brucella melitensis* and *B. abortus* are associated with most of the reported infections in humans in the sub-Saharan Africa. *Brucella* spp. are naturally host-specific but in some circumstances, some strains cause multi-host infections. *B. melitensis* and *B. suis*, for example, cause caprine/ovine and porcine brucellosis and can also infect cattle. Its economic impacts are associated with livestock productivity losses (longer calving intervals, reduced growth, increased incidences of abortion, infertility, and calf mortality) and restrictions on livestock trade [4]. Human infections also prevent infected individuals from engaging in productive occupations.

In livestock, *Brucella* transmission primarily occurs via contact with infected aborted material and ingestion of contaminated feed [5]. Other modes of transmission include natural mating or artificial insemination. Nomadic pastoralism [6] and large herd sizes [7] have been identified as key predictors for exposure in livestock. Humans get exposed to the *Brucella* spp. from animal reservoirs through consumption of unpasteurized dairy products and undercooked meat products, inhalation of contaminated dust and contact with infected animal body fluids or tissues [5]. Person-to-person transmission of the disease is rare; a few such cases have occurred through breastfeeding, trans-placental transmission, blood transfusion and bone marrow transplantation [8]. Herders, livestock owners, and abattoir workers have the highest risk of exposure [9].

The epidemiology of *Brucella* spp. is poorly known. It is generally thought that the disease is endemic among the nomadic communities but the degree of association between livestock and human exposure levels has rarely been determined. A study conducted in Marsabit County, Kenya reported a 6-fold increase in the odds of human seropositivity in household that had a seropositive animal compared to those that did not [10]. This estimate, though, was based on univariable analyses and potential confounding was not accounted for. Another similar study conducted in Togo established that brucellosis was not a major human health problem in the area as very few seropositive individuals were found [11]. The disease was recently ranked among the top five most important zoonotic diseases in the country based on its human health impacts [12]. Part of the challenge arises from the fact that livestock cases, unlike those of humans, often do not manifest any signs apart from initial abortions which occur before immunity is attained, yet infected animals continue to shed the pathogen through milk or genital secretions. It will be difficult therefore to identify carrier animals unless more sensitive and intensive risk-based surveillance measures are deployed.

In this study, we determined patterns of occurrence of *Brucella* spp. seroprevalences in livestock and humans at the household and village levels in Tana River and Garissa Counties, Kenya to obtain data that can inform the development of One Health surveillance measures. The key focus was to determine the strength of association between livestock and human exposures in the rural areas, and to investigate patterns of clustering of the disease at the household and village levels.

## Methods

### Study area

This study was carried out in 2013 – 2014 in Bura and Hola irrigation schemes in Tana River County, and Ijara and Sangailu, Garissa County (Figure 1). The study sites have been described by Bett et al. [13]. Briefly, Bura irrigation and settlement scheme covered 2,100 ha with a tenant population of slightly over 2,000 households settled in 10 villages while Hola irrigation and settlement scheme covered 1,011 ha and had 700 farming households settled in 6 villages. Long-term annual rainfall averaged 460mm and had a trimodal distribution with a major peak in October– December, subsidiary peak in March-May, and minor peak in August-September. The average temperature was 34ºC. Ijara and Sangailu fell under Ijara sub-County which borders Lamu County and Boni forest to the East and Tana River County to the West. Its annual rainfall ranged between 750mm – 1000 mm while the mean temperature ranged between 15°C – 38°C.

**Figure 1:**
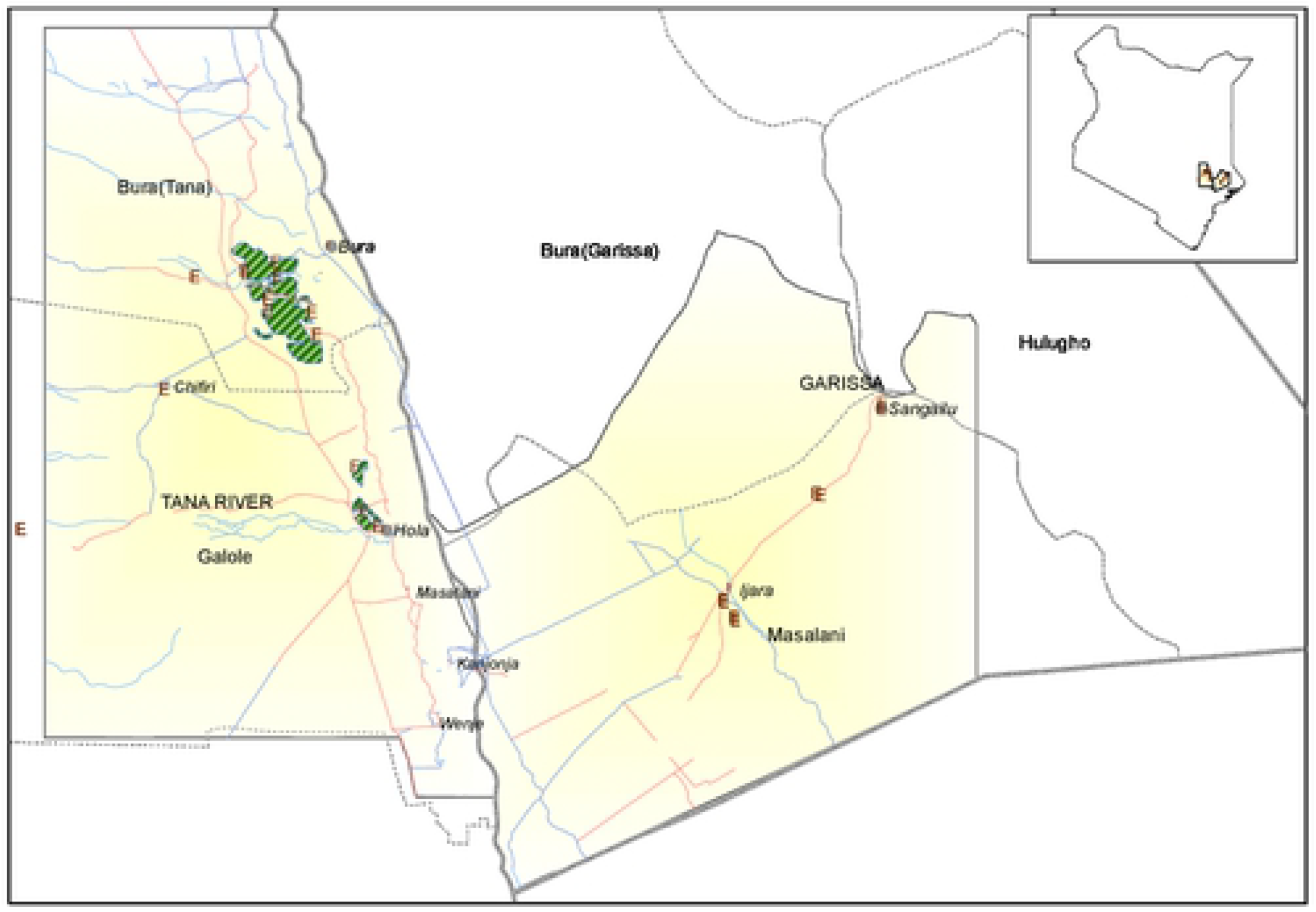
Map showing the study site (ILRI GIS Map, 2013)

### Study design

A cross-sectional study design was used in the study with the primary sampling units being individual livestock and people within selected households. The study aimed to estimate brucella seroprevalence in the target area.

The sample size required was determined using the algorithm described by Humphry et al. [14]. Limited surveys had been done in the area to determine seroprevalences of the disease in livestock and humans, and so *a priori* seroprevalences of 50% were used to obtain the highest possible sample size. Other inputs included desired precision of the test, assumed to be 5%, a confidence level of 95%. The sensitivity and specificity of the SVANOVIR^®^ *Brucella*-Ab C-ELISA that had been identified for use in livestock at the design stage of the study is estimated to be 99.5% and 99.6%, respectively in cattle [15]. A naïve sample size of 400 animals/humans was estimated based on these parameters. Livestock and humans from the same households were however expected to have similar exposure factors and hence their brucella exposures were likely to be correlated. The estimated sample sizes were therefore adjusted for the design effect assuming an intra-household correlation coefficient of 0.20 and that each household had 20 livestock and 5 people. No studies have been done to estimate ICC for brucella but recommendations given by Otte and Gumm [16] suggest that, for most infectious diseases, ICC values range between 0.04 – 0.42, with most values being less than 0.2. Adjusted sample sizes determined were 1920 animals and 720 people. This indicated that 144 households with both livestock and people were required for the study.

A sampling frame comprising a list of households in the study sites was drawn up with the help of the village headmen and the managers of the Bura and Hola irrigation schemes and used in the random selection of households and herds. A household was defined as a group of people who lived together and shared common livelihood activities under a common household head while a herd defined a group of animals owned by a household.

### Animal sampling

Animal sampling was implemented by experienced technicians from the Department of Veterinary Services. Data collected from each animal during blood sampling included age (described as calf, weaner or adult), sex, breed and body condition score. These data were collected using electronic forms designed using Open Data Kit (ODK) application and downloaded to smart phones.

Up to 20 animals were randomly selected in each household/herd for sampling. Selected animals were manually restrained, and blood collected by jugular venipuncture using a vacutainer needle. Plain and EDTA-coated bar-coded vacutainer tubes were each used to collect 5-10ml of blood. Serum was extracted from the plain vacutainer tube at the end of each day at the local livestock office by centrifuging clotted blood at 3,000 rotations per minute for 5 minutes. Extracted serum sample was transferred using sterile Pasteur pipettes into 2ml sterile bar-coded cryovials. Whole blood from EDTA-coated vacutainer tubes was also transferred to 2ml sterile bar-coded cryovials. These samples were stored and transported in dry ice to ILRI Biorepository Unit in Nairobi. Serum and blood samples for each animal were prepared in duplicates.

### Human sampling

A short questionnaire was administered to each subject after the consent process to collect data on age, sex, occupation (farmer, pastoralist, herder, or student) and highest level of education attained. To define occupation, farmers included individuals who grew crops or kept livestock under sedentary production system particularly in the irrigated areas; pastoralists included individuals who kept livestock as a the main source of livelihood and grazed them in communal grazing sites; herders included people that were employed to look after other people’s livestock; and students included household members who were in school or college at the time of sampling. These questionnaires were also administered using ODK forms downloaded to smart phones.

Subjects were prepared for blood sampling by being seated in a comfortable position. The hand used for blood collection was supported and a tourniquet applied above the elbow to distend the veins. The blood collection site was then disinfected using alcohol swabs and an appropriate vacutainer needle (21G for adults and 23G for children) used together with 10ml bar-coded plain and EDTA-coated vacutainer tubes used to collect blood. Each tube was used to collect 5 ml of blood. When adequate blood samples had been obtained, the tourniquet then the needle were removed gently and carefully disposed in the sharps container without recapping and gentle pressure applied to the puncture site for 30 seconds using a cotton swab. The swab was later disposed of and elastoplast^®^ applied to the punctured site. Whole blood from EDTA-coated tubes and serum extracted from plain vacutainer tubes were each transferred to 2ml sterile bar-coded cryovials at the local health centres. These samples were kept and transported in dry ice to biorepository unit at the International Livestock Research Institute (ILRI). Serum and blood samples for each subject were prepared in duplicates.

### Questionnaire survey

Semi-structured questionnaires (also designed using the ODK application) were administered to the household heads. This collected information on participants’ demographics, knowledge, attitudes and practices of the communities in relation to brucellosis transmission and control as well as information on possible risk factors. Sampling forms were used to collect further information on livestock ownership, herd size, type of livestock owned, age, sex and occupation.

### Laboratory screening

Human samples were tested using anti-*Brucella* IgG enzyme-linked immunosorbent assay (ELISA) (Demeditec Diagnostics GmbH Kiel, Germany) while animal samples were tested using *Brucella* competitive ELISA kit (SVANOVIR^®^ *Brucella*-Ab C-ELISA). The anti-*Brucella* IgG ELISA is designed for qualitative measurement of IgG class of antibodies in human plasma or serum. It has a high sensitivity (>99%) but it is vulnerable to non-specific reactions particularly to cross-reactions with bacteria having lipopolysaccharides (LPS). The C-ELISA kit is based on S-LPS antigen and it can detect *B. abortus, B. melitensis* and *B. suis*. This kit is more specific but less sensitive compared to the anti-*Brucella* IgG ELISA since it targets specific epitopes of the *Brucella* LPS not shared with other pathogens. All assays were performed as stipulated in the manufacturer’s standard operating procedures.

### Data management and analysis

Data collected were uploaded to the ILRI server at the end of each day using internet-enabled smart phones. Each data file was saved as a comma delimited file (.csv). They were exported to Microsoft Access 2010 for cleaning and merging, and finally to STATA version 13 for analysis.

Both livestock and human data were subjected to similar descriptive and inferential statistics. Descriptive analyses commenced with the determination of the overall frequencies and prevalence (with 95% confidence interval) of seropositive subjects in animals and humans. These estimates were further stratified by all the categorical variables. For livestock data, categorical variables included species (cattle, sheep or goats), sex, age (calf/lamb/kid, weaner or adult), area (Bura or Hola) and land use (pastoralism or irrigation), while for human data these were sex, age (≤17 years, 18-40 years or >40 years), occupation (pastoralism, farmer, student, other [e.g. business, housewife, chief, etc.]), location (Ijara, Sangailu, Bura and Hola) and land use (pastoralism or irrigation). Occupation was collapsed into the four levels defined above because the original form of the variable had up to 13 levels with some of them having sparse data. Age (from the human data) was also recoded into a 3-level categorical variable because the original form, which was captured as a continuous variable, did not satisfy the linearity assumption during modelling (described below). To determine the relationship between seropositivity in livestock and humans at the household level, a dummy variable was created from livestock seropositivity data (from Tana River County) to indicate whether there was at least one seropositive animal in a given household (value = 1) or not (value = 0). This was then merged with the human data using the household ID as the primary key.

Univariable logistic regression models were fitted to each dataset to identify unconditional association between each of these variables with their respective outcomes (*Brucella* spp. exposure status in animals and humans). Multivariable random effects logistic regression models were fitted to these data through a combination of backward-forward variable selection technique with α=0.05. The analysis used *melogit* command in STATA, with the default integration method and points (i.e., mean-variance adaptive Gauss–Hermite quadrature and 7, respectively). The analyses commenced with a saturated model that was systematically reduced by removing variables (both fixed and random effects) that had a p value > 0.05 based on likelihood ratio test (lrtest). The ICCs were generated using the command *estat icc*. Deviance residuals and fitted values were generated and used to evaluate the final models developed.

### Herd/household level seroprevalence

Secondary descriptive analyses were done to determine herd and household level seroprevalences of *Brucella* spp.

### Ethics statement

Protocols used for human sampling were reviewed and approved by AMREF Ethics and Scientific Review Committee (with a reference number: P65/2013). On the day of sampling, selected subjects were taken through the consent process by a local clinician before being sampled in presence of a witness identified by the subject. They also gave a written consent – one copy of the signed consent form was kept by the clinician while another was kept by the subject. If the subject was a child aged 5-12 years, only the parent’s consent was obtained. If the selected individual was a child between 13-17 years of age, the subject’s assent was taken together with that of the parent or guardian. For adults (18 years or more), personal consents were required. All these processes required the presence of a witness who verified that adequate information on the research was provided and subjects participated in the research voluntarily.

For the livestock component, protocols were reviewed and approved by the International Livestock Research Institute’s (ILRI) Institutional Animal Care and Uses Committee (IACUC) (reference number 2014.02). ILRI IACUC is registered in Kenya and complies with the UK’s Animals (Scientific Procedures) Act 1986 (http://www.homeoffice.gov.uk/science-research/animal-research/) that contains guidelines and codes of practice for the housing and care of animals used in scientific procedures. The study adhered to the IACUC’s 3R principles of (i) replacement of animal with non-animal techniques, (ii) reduction in the number of animals used, and (iii) refinement of techniques and procedures that reduce pain and distress. Animal owners provided oral consent for animal sampling.

## Results

### Descriptive analyses

A total of 2,025 animals comprising 441 cattle, 961 goats and 623 sheep were sampled from 143 households in 20 villages in Tana River County. Most of these animals (76.59%, n = 1,551) were from villages in the irrigated areas both in Bura and Hola while the rest were from the neighboring pastoral areas. Overall, 71.11% (n = 1,440) of the animals were from Hola. Regarding species distribution, all the sheep and cattle sampled in Bura were black head Persian and Orma Boran breeds, respectively, while a majority (over 99%) the goats were Galla or Galla crosses. A similar pattern was observed in Hola where 91.83% (n = 674) of the goats sampled were Galla breed or their crosses and a large proportion of cattle and sheep sampled were Orma Boran and black head Persian breeds respectively. The mean goat herd size was 29.41 (range of 0 to 120) while that of cattle herds and sheep flocks was 24.11 (range 0 to 200, median 2) and 31.42 (range 0 to 200) respectively. It was not possible to sample livestock in Ijara and Sangailu (Garissa County) due to insecurity challenges.

A total of 1,022 people from 364 households (99 in Ijara, 93 in Sangailu, 100 in Bura and 72 in Hola) were sampled. The number sampled per household ranged from one to six with a mean of 2.81 (SE 0.07). A higher percentage of these subjects, 59.55% (n = 608), were females. Common occupations included pastoralism (35.57%, n = 234), farmers (29.57%, n = 194), and student (19.51%, n = 128). Others were formal employment, herdsman and housewife. The distribution of the subjects by sites was balanced; 30.14% (n=308) were from Bura, 21.82% (n = 223) were from Hola, 23.68% (n = 242) were from Sangailu and the rest 24.36% (n = 249) were from Ijara. In the irrigated area, the source of water was mainly from canals while in the non-irrigated/pastoral area, the main source of water was boreholes and dams.

### *Brucella* spp. seroprevalence

#### Livestock

Seventy out of 2,025 animals (3.41%; 95% C: 2.62 – 4.20%) tested positive for *Brucella* spp. antibodies. The overall seroprevalence in Hola (2.64%, 95% CI: 1.81 – 3.47%) was significantly lower than that in Bura (5.29%, 95% CI: 3.48 – 7.12). The seroprevalences in cattle, goats and sheep were 6.35% (95% CI: 4.06 – 8.63%), 3.33% (2.19 – 4.47%) and 1.44% (0.51 – 2.38%). These estimates were stratified by sex, age, area and land use (pastoral verses irrigation and the results are given in Table 1.

**Table 1:** *Brucella* spp. seroprevalence in livestock in Tana River County, Kenya

| Variable | Levels | Livestock species |  |  |  |  |  |  |  |  |  |  |  |
| --- | --- | --- | --- | --- | --- | --- | --- | --- | --- | --- | --- | --- | --- |
|  |  | Cattle |  |  |  | Goats |  |  |  | Sheep |  |  |  |
|  |  | *n* | Seroprevalence |  |  | *n* | Seroprevalence |  |  | *n* | Seroprevalence |  |  |
| | | | *p* (%) | 95% CI | P<$\chi^2$ | | *p* (%) | 95% CI | P<$\chi^2$ | | *p* (%) | 95% CI | P<$\chi^2$ |
| Sex | Male | 122 | 1.64 | 0.20 – 5.80 | 0.02 | 201 | 3.98 | 1.73 – 7.69 | 0.56 | 143 | 1.39 | 0.16 – 4.96 | 0.95 |
|  | Female | 338 | 7.99 | 5.33 – 11.41 |  | 760 | 3.16 | 2.03 – 4.66 |  | 480 | 1.46 | 0.59 – 2.98 |  |
| Age | Calf/kid/lamb | 77 | 0 | - | 0.00 | 37 | 5.41 | 0.67 – 18.19 | 0.04 | 25 | 4.00 | 0.10 – 20.35 | 0.25 |
|  | Weaner | 147 | 2.04 | 0.41 – 5.85 |  | 146 | 0 | - |  | 105 | 0 | - |  |
|  | Adult | 217 | 11.52 | 7.60 – 16.54 |  | 755 | 3.97 | 2.69 – 5.62 |  | 489 | 1.64 | 0.71 – 3.19 |  |
| Area | Bura | 249 | 6.02 | 3.41 – 9.74 | 0.75 | 198 | 6.06 | 3.17 – 10.35 | 0.02 | 138 | 2.89 | 0.79 – 7.26 | 0.10 |
|  | Hola | 192 | 6.77 | 3.65 – 11.30 |  | 734 | 2.62 | 1.61 – 4.02 |  | 494 | 1.03 | 0.34 – 2.39 |  |
| Land use | Pastoral | 212 | 6.73 | 3.72 – 11.04 | 0.76 | 177 | 6.11 | 3.09 – 10.67 | 0.02 | 86 | 0 |  | 0.23 |
|  | Irrigation | 230 | 6.00 | 3.32 – 9.88 |  | 781 | 2.69 | 1.67 – 4.08 |  | 537 | 1.68 | 0.77 – 3.16 |  |

*n* – Total number of animals sampled
*p* – seroprevalence
CI – confidence interval

#### Humans

Using all the data from the four sites, 366 subjects (35.81%, 95% CI: 32.87 – 38.84%) were seropositive for *Brucella* spp. Table 2 gives the distribution of these cases by the five independent factors considered. The table also show similar results with reduced data set from two sites in Tana River County. In both cases, males had higher prevalence than females and age was also positively associated with increased levels of exposure. Regarding occupation, pastoralists had higher seroprevalence than farmers, students or other occupations (including business, formal employment, etc). The results further show that of the four areas sampled, Sangailu had the highest seroprevalence of 54.13%, followed by Ijara, Hola and Bura with 47.39%, 26.46 and 18.83% seroprevalences, respectively.

**Table 2.**
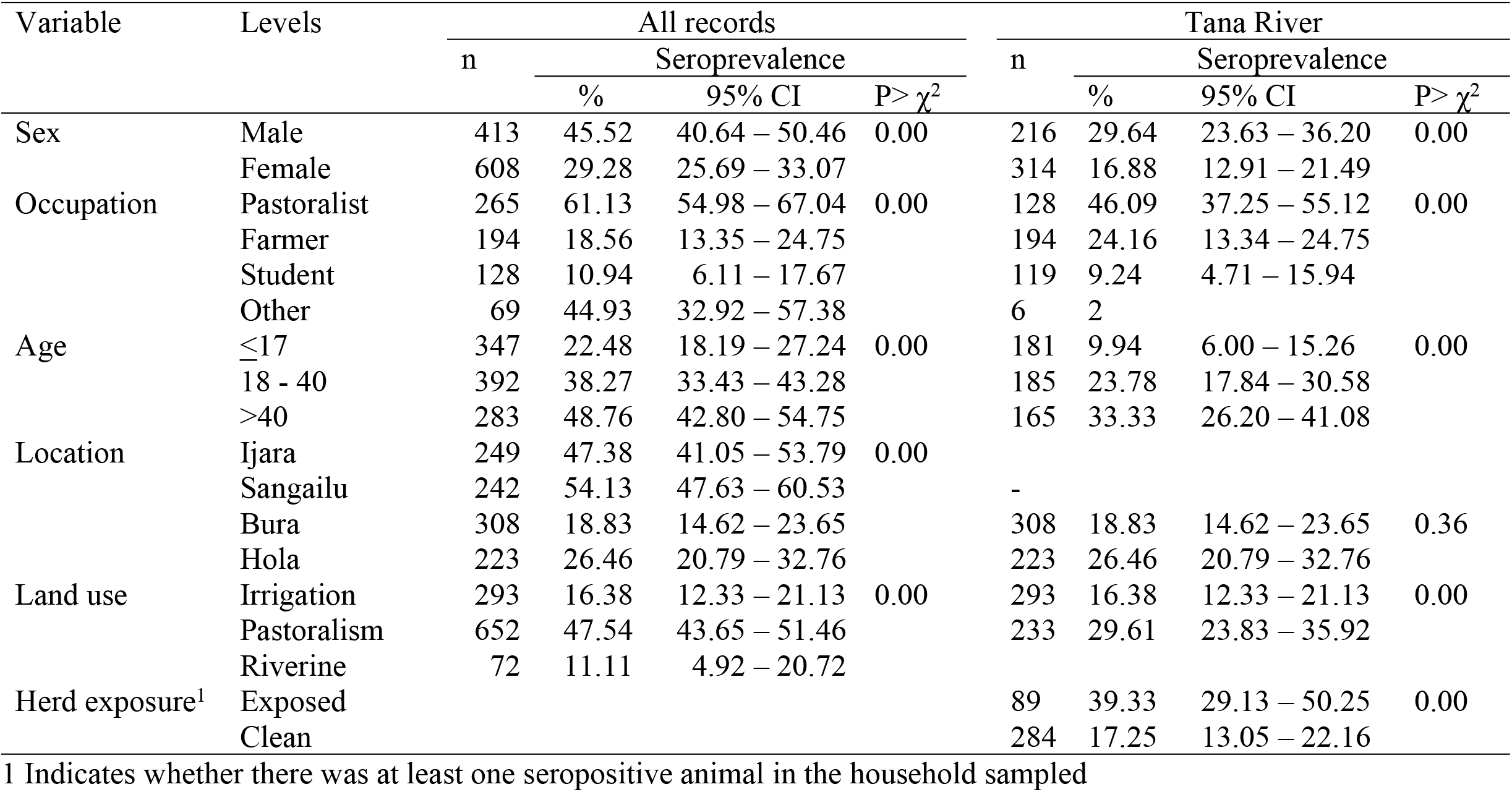
*Brucella* spp. seroprevalence in humans presented separately first for all the data from Garissa and Tana River counties, and secondly for records from Tana River County (Bura and Hola areas) where livestock were also sampled

### Risk factor analysis

#### Univariable analysis

Univariable analyses on the livestock data identified five out of six variables as being significant in the naïve logistic regression model used. Age of an animal, species (cattle, sheep, goats), area from which it came from (Hola verses Bura), and land use type in the area (irrigation verses pastoralism) were significant as fixed effects while household ID and village ID were significant as random effects. Sex was not significant (p= 0.23) and so this variable was not used in the subsequent multivariable analyses. Herd size at the household level was also significant. This variable was used as a continuous variable since it satisfied conditions for the linearity assumption.

Regarding human data, all the independent variables listed in Table 2 were significant in univariable analyses. An additional variable (family size) was included in this analysis but it was found to be non-significant. It was therefore not considered for the subsequent multivariable analyses. Comparable results were obtained for the two sets of data described in Table 2 (i.e., with all the data from Garissa and Tana River counties versus those from Tana River County alone where livestock sampling was done). Multivariable analysis on human data therefore used data from Tana River only since these allowed analyses on the association between livestock and human seroprevalences to be evaluated.

#### Multivariable analysis

Table 3 gives the results of final mixed effects logistic regression model fitted to the livestock data. Two significant fixed effects variables were age of the animal and area (though this is barely significant at 95% confidence). We found an ordinal increase in the risk of exposure with age, while for area, an animal from Bura was 3.73 times more likely to be exposed to *Brucella* spp. than that from Hola. Land use and herd size were not significant in this model. A model used to investigate interactions between age and area did not converge.

**Table 3.** Output from a mixed effects logistic regression model showing the association between *Brucellosis* spp. seropositivity in livestock and risk factors studied

| Variables | Levels | Odds Ratio |  |  | Z | P>Z |
| --- | --- | --- | --- | --- | --- | --- |
|  |  | Estimate | SE | 95% CI |  |  |
| *Fixed effects* |  |  |  |  |  |  |
| Age | Kid/lamb/calf | 0.27 | 0.17 | 0.08 – 0.92 | -2.08 | 0.04 |
|  | Weaner | 0.13 | 0.08 | 0.04 – 0.43 | -3.35 | 0.01 |
|  | Adult | 1.00 |  |  |  |  |
| Area | Hola | 0.24 | 0.17 | 0.06 – 0.94 | -2.04 | 0.04 |
|  | Bura | 1.00 |  |  |  |  |
| Constant |  | 0.06 | 0.03 | 0.02 – 0.17 | -5.16 | 0.00 |
| *Random effects* |  |  |  |  |  |  |
| Herd ID/Village |  | 0.87 | 0.45 | 0.31 – 2.43 |  |  |
| Village ID |  | 1.30 | 0.81 | 0.38 – 4.39 |  |  |

Log likelihood -267.31, number of observations 1,998

Village and herd IDs (with herd IDs nested within village) were used in the model as random effects variables and both were significant (p = 0.00) in the model. The variances (SE) associated with observations within a household and village were 1.30 (SE 0.81) and 0.87 (SE 0.45), respectively. The internal correlation coefficient of exposures between animal within households was 0.40 (95% CI: 0.22 – 0.60) and between animals from different households within a village was 0.24 (95% CI: 0.08 – 0.52). Deviance residuals ranged between −1 and 3.

Outputs from the model fitted to the human data (Tana River county) is presented in Table 4. Four variables – age, gender, land use and exposure status of livestock in the household were significant. There was an ordinal increase in the odds of seropositivity with age, and males had higher odds of exposure than females. Similarly, people who lived in pastoral areas had higher odd of exposure. The main finding was that the odds of exposure were 3.35 times higher in people from households that had at least one seropositive animal compared to those that did not. The inclusion of herd exposure at the household level rendered the village random effects variable to be insignificant (p = 0.13). The final model therefore had only one random effects variable – household, with an ICC estimates of 0.21 (95% CI: 0.06 – 0.52).

**Table 4.** Results from the random effects logistic regression model fitted to data on human *Brucella* spp. exposure in Tana River County, Kenya

| Variable | Level | Odds Ratio |  |  | z | P>z |
| --- | --- | --- | --- | --- | --- | --- |
|  |  | Estimate | SE | 95% CI |  |  |
| *Fixed effects* |  |  |  |  |  |  |
| Age | ≤17 | 0.13 | 0.07 | 0.05 – 0.36 | -3.91 | 0.00 |
|  | 18 - 40 | 1.00 |  |  |  |  |
|  | >40 | 2.16 | 0.77 | 1.08 – 4.34 | 2.17 | 0.03 |
| Gender | Male | 2.33 | 0.78 | 1.20 – 4.50 | 2.51 | 0.01 |
|  | Female | 1.00 |  |  |  |  |
| Land use | Irrigation | 0.18 | 0.08 | 0.08 – 0.42 | -3.97 | 0.00 |
|  | Pastoral | 1.00 |  |  |  |  |
| Herd exposure | Exposed | 3.35 | 1.35 | 1.51 – 7.41 | 2.98 | 0.00 |
|  | Clean | 1.00 |  |  |  |  |
| Constant |  | 0.44 | 0.18 | 0.19 – 0.98 | -2.00 | 0.05 |
| *Random effect* |  |  |  |  |  |  |
| Household ID |  | 0.86 | 0.62 | 0.21 – 3.56 |  |  |

Log likelihood -150.81, number of observations 364

Residual analyses conducted for both models indicated that there were no outliers as deviance residuals ranged between −2 and 2.5.

### Herd/household level *Brucella* spp. seroprevalence

The overall herd level *Brucella* spp. seroprevalence was 25.87% (95% CI: 18.61 – 33.14%). This was positively associated with herd size, and herds from Hola had higher odds of exposure than those from Bura (Table 5). Herd size was kept in the model as a continuous variable since it met the linearity assumption. The internal correlation coefficient indicating the correlation of observations between herds from the same village was estimated to be 0.34 (95% CI: 0.10 – 0.69).

**Table 5.** Factors affecting herd-level *Brucella* spp. seroprevalence in livestock in Tana River County

| Variable | Level | Odds Ratio |  |  | z | P>z |
| --- | --- | --- | --- | --- | --- | --- |
|  |  | Estimate | SE | 95% CI |  |  |
| *Fixed effects* |  |  |  |  |  |  |
| Herd size<sup>1</sup> |  | 1.01 | 0.004 | 1.01 – 1.02 | 3.00 | 0.00 |
| Area | Bura | 8.82 | 7.98 | 1.50 – 51.93 | 2.41 | 0.02 |
|  | Hola | 1.00 |  |  |  |  |
| Constant |  | 0.06 | 0.04 | 0.02 – 0.21 | -4.41 | 0.00 |
| *Random effect* |  |  |  |  |  |  |
| Village ID |  | 1.67 | 1.28 | 0.37 – 7.46 |  |  |

Log likelihood -62.38, number of observations 143

The overall household-level seroprevalences was 60.16 (95% CI: 54.93 – 65.23%). This varied significantly by location. Sangailu and Ijara had the highest household-level seroprevalences of 81.72% (95% CI: 72.35 – 88.98) and 68.67% (95% CI: 58.59 – 77.64%), respectively. Those for Hola and Bura were 51.39% (95% CI: 39.31 – 63.35%) and 38.00% (95% CI: 28.48 – 48.25%), respectively.

## Discussion

We determined *Brucella* spp. seroprevalences and their risk factors in livestock and humans in two contrasting ecological regions that practiced contrasting livelihood practices (i.e., pastoralism and crop irrigation). The subject-level seroprevalences observed (ranging between 2.67 – 5.39% in livestock and 18.83 – 54.13% in humans) are similar to those that have been reported previously in other parts of Kenya. A study conducted by Osoro et al. [10] in Kiambu, Kajiado and Marsabit Counties provided *Brucella* spp. seroprevalences in livestock of 1.2 – 13.5% and in people of 2.4% – 46.5%. The herd and household level seroprevalences obtained in the current study were also similar to those reported earlier by Osoro et al.[10]. In this study, however, the household seroprevalences (of 81.72% (95% CI: 72.35 – 88.98) observed in Sangailu where were exceptionally higher than any of those reported earlier. This area (with Ijara) had a large population of livestock and wildlife and the local people practiced pastoralism (discussed below) which could have promoted frequent exposure to *Brucella* spp. and other zoonotic pathogens.

The multivariable models we used identified three factors – age, area and herd size – as being significant predictors of *Brucella* spp. exposure in livestock. Older animals were generally associated with increased risk of *Brucella* spp. exposure compared to younger ones. This is a common finding with several possible explanations. First, compared to the young, older animals have had more chances of encountering infectious hosts, and given that *Brucella* spp. antibodies can persist in body tissues for a long time, seropositivity in adults might be representing past cumulative exposures. Secondly, the communities involved in the study kept breeding herds for many years unlike young animals that could be sold whenever there was need for money. We therefore expect a higher replacement rate of the younger, non-breeding animals in a herd which might result in the replacement of the few exposed subjects.

We observed a positive association between herd size and *Brucella* spp. seroprevalence. This relationship has been reported in Ethiopia [17], Zambia [18], and Pakistan [19] and many other areas. Large herds, especially those that comprise mixed species allow more efficient transmission of *Brucella* spp. [20]. This is because large herds have a high number of susceptible hosts and higher numbers of effective contacts, permitting endemic circulation of the pathogen over time. In pastoral areas, large herds are often raised under traditional management practices that further contribute to *Brucella* spp. transmission. These include communal grazing, natural breeding, and confinement in small enclosures that encourage close contact, especially in the night. *Brucella* spp. transmission models show that the disease is persistent in pastoral areas and its interventions should include those that manage ecosystems such as land reform, maintenance of adequate stocking rates, and an integrated social and economic development [20].

The intra-herd correlation coefficient of *Brucella* spp seropositivity estimated in the study (of 0.39) was about two times higher than that between herds (0.18), an indication that livestock cases clustered more at the herd than at the village level. This finding suggests that within-herd exposures played a stronger role than those between herds. Pasture contamination is considered as one of the main modes of transmission of the pathogen in livestock but the duration over which the pathogen remains viable in the environment in this area, given the high day-time temperatures, is not known. At the same time, livestock share grazing and watering areas which further increases chances of pathogen transmission. Based on observations made by Otte and Gumm [16] that most infectious diseases have ICC values of less than 0.2, it is apparent that brucellosis is one of the diseases with the highest inter-herd correlation coefficient.

Many studies have identified risk factors for *Brucella* spp exposure in humans. This study identified age, gender and land use as being important determinants of *Brucella* spp. exposure. Older people had higher odds of exposure than younger, and men had higher odds of exposure than women. A study conducted in the northern Kenya indicated that regular ingestion of raw milk, herding, milking and feeding goats and handling of hides were associated with increased of exposure to *Brucella* spp. [10]. In Northern Tanzania, Kunda et al [21] also confirmed that assisting animals give birth enhances the risk of exposure to *Brucella* spp. In general, any livelihood practices that encourages intense contact with infected animals, or consumption of its products would increase the risk of brucellosis.

On the relationship between human and animal *Brucella* spp. seroprevalences, this study established that controlling for the other covariates, the odds of a person being exposed to *Brucella* spp. was 3.34 times higher in households that had at least one seropositive animal compared to people in households that did not. Osoro et al. [10] conducted a similar analysis in northern Kenya but they used univariable models which can be more sensitive to confounding than those used here. Our observations on high degree of clustering at the household/herd level and a strong association between human and livestock exposures suggest an important opportunity for the deployment of One Health surveillance and control for the disease. It is possible to use initial cases identified in animals or humans to identify households where additional cases are likely to be occur. This requires exchange of surveillance data between public and animal health institutions, and the implementation of coordinated responses to manage the existing cases and prevent further exposures. Public health workers, for example, should be able to advice animal health workers to trace infected animals in households where a patient diagnosed with brucellosis comes from to reduce chances of re-infection, and to protect other members of the household. For those cases where the animal health workers cannot access screening materials, participatory techniques that include using case definitions that includes history of abortion should be used to detect and remove infected animals.

The study had a few limitations. First, it was designed as a cross sectional survey which can’t investigate dynamic changes in *Brucella* spp seroprevalence. Secondly, samples were screened using serological tests which do not (i) allow for the characterization of the pathogen by species as it targets immunodormant *Brucella* antigens associated with the smooth LPS that is shared by multiple naturally occurring biovars of *B. abortus*, *B. melitensis*, *B. suis* and others [19], (ii) allow for confirmation of a subject’s infection status as the presence of antibodies does not necessarily suggest current infection. Humoral IgG responses remain detectable for a long time (and might last for years) and the excretion of *Brucella* spp. occurs occasionally, more so during abortions [19]. The isolation of the bacteria and the detection of its DNA by PCR, and to some extent the detection of IgM antibodies, remain the most reliable means of detecting active infections.

## Conclusions

Our study gives evidence of a strong association between human and animal seropositivity at household level. The other important risk factors are occupation, age, sex, and land use; these should be taken into consideration when designing interventions against the disease. There is also high prevalence of brucellosis in human and livestock, and it is higher in the predominantly pastoralist communities. We recommend further research to characterize *Brucella* spp. strains that are circulating in these areas to understand how these pathogens are shared between wildlife, livestock and people.

## Acknowledgements

We thank all the people who participated in the study including livestock owners, veterinarians from Tana River and Garissa Counties, clinicians from Bura, Hola, Ijara and Sangailu health centres, and the local administrative officers from these areas. The serosurveillance data were collected by John Muriuki and Damaris Mwololo.

## Supporting information

S1 Checklist: STROBE Checklist

## References

1. Corbel MJ. Brucellosis: an overview. Emerg Infect Dis. 1997;3: 213–221. doi:10.3201/eid0302.970219

2. Callaghan DO, Whatmore AM. Brucella genomics as we enter the multi-genome era. 2011;10: 334–341. doi:10.1093/bfgp/elr026

3. Roth F, Zinsstag J, Orkhon D, Chimed-Ochir G, Hutton G, Cosivi O, et al. Human health benefits from livestock vaccination for brucellosis:case study. Bull World Health Organ. 2003;81: 867–876. doi:10.1590/S0042-96862003001200005

4. McDermott J, Grace D, Zinsstag J. Economics of brucellosis impact and control in low-income countries. Rev Sci Tech. 2013;32: 249–261.

5. Corbel MJ. Brucellosis in humans and animals. WHO. 2006; 1–102. doi:10.2105/AJPH.30.3.299

6. Njeru J, Wareth G, Melzer F, Henning K, Pletz MW, Heller R, et al. Systematic review of brucellosis in Kenya: disease frequency in humans and animals and risk factors for human infection. BMC Public Health. England; 2016;16: 853. doi:10.1186/s12889-016-3532-9

7. McDermott JJ, Arimi SM. Brucellosis in sub-Saharan Africa: epidemiology, control and impact. Vet Microbiol. Netherlands; 2002;90: 111–134.

8. Tuon FF, Gondolfo RB, Cerchiari N. Human-to-human transmission of Brucella – a systematic review. Trop Med Int Heal. 2017;22: 539–546. doi:10.1111/tmi.12856

9. de Glanville WA, Conde-Álvarez R, Moriyón I, Njeru J, Díaz R, Cook EAJ, et al. Poor performance of the rapid test for human brucellosis in health facilities in Kenya. PLoS Negl Trop Dis. 2017;11: e0005508. doi:10.1371/journal.pntd.0005508

10. Osoro EM, Munyua P, Omulo S, Ogola E, Ade F, Mbatha P, et al. Strong Association Between Human and Animal Brucella Seropositivity in a Linked Study in Kenya, 2012-2013. Am J Trop Med Hyg. 2015;93: 224–31. doi:10.4269/ajtmh.15-0113

11. Dean AS, Bonfoh B, Kulo AE, Boukaya GA, Amidou M, Hattendorf J, et al. Epidemiology of Brucellosis and Q Fever in Linked Human and Animal Populations in Northern Togo. PLoS One. 2013;8: 1–8. doi:10.1371/journal.pone.0071501

12. Munyua P, Bitek A, Osoro E, Pieracci EG, Muema J, Mwatondo A, et al. Prioritization of Zoonotic Diseases in Kenya, 2015. PLoS One. United States; 2016;11: e0161576. doi:10.1371/journal.pone.0161576

13. Bett BK, Said MY, Sang R, Bukachi S, Wanyoike S, Kifugo SC, et al. Effects of flood irrigation on the risk of selected zoonotic pathogens in an arid and semi-arid area in the eastern Kenya. PLoS One. 2017;12: 1–15. doi:10.1371/journal.pone.0172626

14. Humphry RW, Cameron A, Gunn GJ. A practical approach to calculate sample size for herd prevalence surveys. Prev Vet Med. 2004;65: 173–188. doi:10.1016/j.prevetmed.2004.07.003

15. Svanova. The best way to detect brucellosis in livestock herds [Internet]. 2019 [cited 16 May 2019]. Available: https://www.svanova.com/content/dam/internet/ah/svanova/dk_EN/documents/porcine/Brucella-C_Infosheet_02.pdf

16. Otte MJ. Intra-cluster correlation coefficients of 20 infections calculated from the results of cluster-sample surveys. Prev Vet Med. 1997;5877.

17. Terefe Y, Girma S, Mekonnen N, Asrade B. Brucellosis and associated risk factors in dairy cattle of eastern Ethiopia. Trop Anim Health Prod. 2017;49: 599–606. doi:10.1007/s11250-017-1242-7

18. Muma JB, Samui KL, Oloya J, Munyeme M, Skjerve E. Risk factors for brucellosis in indigenous cattle reared in livestock–wildlife interface areas of Zambia. Prev Vet Med. 2007;80: 306–317. doi:10.1016/j.prevetmed.2007.03.003

19. Ali S, Akhter S, Neubauer H, Melzer F, Khan I, Abatih EN, et al. Seroprevalence and risk factors associated with bovine brucellosis in the Potohar Plateau, Pakistan. BMC Res Notes. 2017;10: 73. doi:10.1186/s13104-017-2394-2

20. Racloz V, Schelling E, Chitnis N, Roth F. Persistence of brucellosis in pastoral systems. Sci Tech Rev Off Int des Epizoot. 2013;32: 61–70. Available: http://europepmc.org/abstract/MED/23837365%5Cnpapers3://publication/uuid/1C112554-B814-4CAA-B1ED-97117DC8E7F3

21. Kunda J, Fitzpatrick J, French N, Kazwala R, Kambarage D, Mfinanga GS, et al. Quantifying Risk Factors for Human Brucellosis in Rural Northern Tanzania. Noor AM, editor. PLoS One. 2010;5: e9968. doi:10.1371/journal.pone.0009968

